# Early induction of SARS-CoV-2 specific T cells associates with rapid viral clearance and mild disease in COVID-19 patients

**DOI:** 10.1101/2020.10.15.341958

**Authors:** Anthony T. Tan, Martin Linster, Chee Wah Tan, Nina Le Bert, Wan Ni Chia, Kamini Kunasegaran, Yan Zhuang, Christine Y. L. Tham, Adeline Chia, Gavin J. Smith, Barnaby Young, Shirin Kalimuddin, Jenny G. H. Low, David Lye, Lin-Fa Wang, Antonio Bertoletti

**Affiliations:** Programme in Emerging Infectious Diseases, Duke-National University of Singapore Medical School, Singapore; National Centre for Infectious Diseases, Singapore; Department of Infectious Diseases, Tan Tock Seng Hospital, Singapore; Lee Kong Cian School of Medicine, Singapore; Department of Infectious Diseases, Singapore General Hospital, Singapore; Yong Loo Lin School of Medicine, National University Singapore, Singapore; Singapore Immunology Network, A*STAR, Singapore

## Abstract

Virus-specific humoral and cellular immunity act synergistically to protect the host from viral infection. We interrogated the dynamic changes of virological and immunological parameters in 12 patients with symptomatic acute SARS-CoV-2 infection from disease onset to convalescence or death. We quantified SARS-CoV-2 viral RNA in the respiratory tract in parallel with antibodies and circulating T cells specific for various structural (NP, M, ORF3a and spike) and non-structural proteins (ORF7/8, NSP7 and NSP13). We observed that while rapid induction and quantity of humoral responses were associated with increased disease severity, an early induction of SARS-CoV-2 specific T cells was present in patients with mild disease and accelerated viral clearance. These findings provide further support for a protective role of SARS-CoV-2 specific T cells over antibodies during SARS-CoV-2 infection with important implications in vaccine design and immune-monitoring.

## Introduction

In December 2019, a new coronavirus was detected in Wuhan, China in several patients with pneumonia and was later named SARS-CoV-2(Zhou et al., 2020). The illness (COVID-19) resulting from SARS-CoV-2 infection is reported to be multi-faceted with inflammation of the respiratory tract causing the leading symptoms of fever and dry cough {Chen:2020kl}. Both humoral (Long et al., 2020a) (Gudbjartsson et al., 2020; Sun et al., 2020) and cellular (Grifoni et al., 2020) (Braun et al., 2020; Le Bert et al., 2020) (Weiskopf et al., 2020) components of adaptive immunity are induced in SARS-CoV-2 infected individuals but their roles in viral control or disease pathogenesis still needs to be clarified. Viral clearance and reduced disease severity have been associated with a coordinated activation of humoral and cellular anti-viral immunity (Rydyznski Moderbacher et al., 2020) and robust T cell responses (Takahashi et al., 2020). On the other hand, a positive relation between the magnitude of SARS-CoV-2 antibodies (Hung et al., 2020; Long et al., 2020b) or T cells(Peng et al., 2020) with disease severity have also been reported. However, most of these studies have analysed patients during the convalescent phase of infection and only a single study reported the dynamic changes of viral and immunological parameters in severe COVID-19 patients during the initial phases of infection (Weiskopf et al., 2020). To fill this gap, we longitudinally followed-up twelve patients with SARS-CoV-2 infection from symptom onset to convalescence or death. We quantified SARS-CoV-2 viral load in the upper respiratory tract and virus-specific antibodies and T cells at multiple time points. Our data reveal a direct association between early induction of SARS-CoV-2 specific T cells and rapid control of viral infection.

## Results

### Dynamics of SARS-CoV-2 replication

Relative quantities of SARS-CoV-2 in the respiratory tract and its persistence in each individual patient was calculated by utilizing the number of RT-PCR cycles as a proxy of viral quantity (Figure 1). Duration of infection was defined from symptom onset till RT-PCR negativity (two negative SARS-CoV-2 RT-PCR 24 hours apart). The patients displayed different profiles of virus quantity and persistence: short duration of infection was detected in four patients (P18, 04, 02 and 15) who became RT-PCR negative within 15 days of symptom onset. Five (P11, 16, 06, 09 and 03) were RT-PCR negative at around day 17-24, while two (P10 and 12) became RT-PCR negative almost 1 month after onset of symptoms. One patient (P05) who succumbed to infection was persistently SARS-CoV-2 RT-PCR positive till day 31 post symptom onset when he demised. Peak viral quantities was also positively correlated with the duration of infection (Figure 1 insert). In addition, while patients who eliminated the virus within 15 days experienced mild respiratory symptoms (presence of fever or respiratory symptoms but not requiring supplemental oxygen), patients with moderate (requiring oxygen supplementation FiO2<0.5) and severe (requiring oxygen supplementation FiO2>0.5, high flow oxygen and/or mechanical ventilation) symptoms eliminated the virus from upper respiratory tract later.

**Figure 1.**
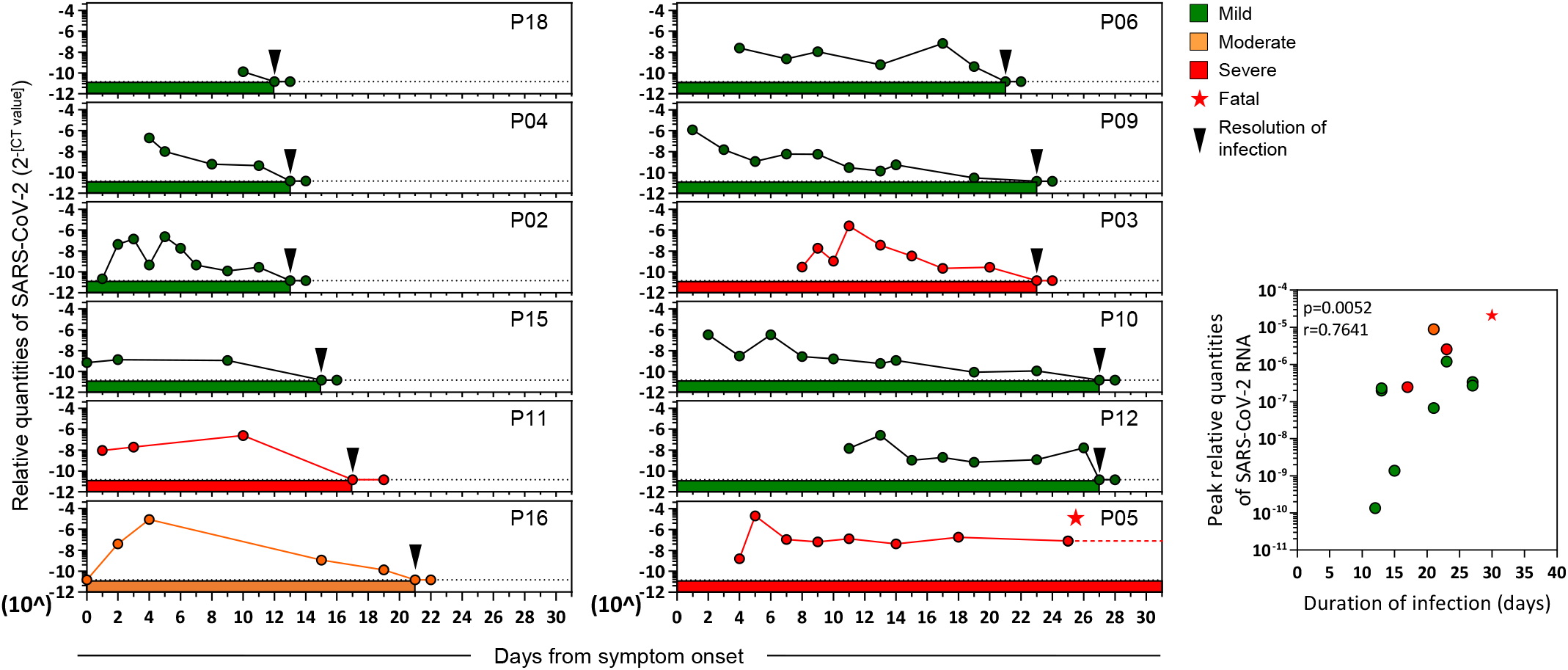
Relative quantities of SARS-CoV-2 in the upper respiratory tract of symptomatic COVID-19 patients during acute infection. Longitudinal RT-PCR quantification of SARS-CoV-2 RNA in the upper respiratory tract of COVID-19 patients (n=12) with variable symptom severity from symptom onset till RT-PCR negativity. Dotted lines denote the positive cut-off. Insert shows the correlation between the peak relative quantities of SARS-CoV-2 and the duration of infection.

### Dynamics of SARS-CoV-2 specific antibody response

We then characterized the kinetics of anti-SARS-CoV-2 antibody appearance. Virus neutralization ability was tested longitudinally utilizing the surrogate virus neutralization test (sVNT) that quantified the ability of serum antibodies to inhibit the binding of Spike RBD to ACE2 receptor *in vitro* (Tan et al., 2020). We also quantified anti-RBD, anti-S1 (S1 domain of spike) and anti-NP (nucleoprotein) IgG and IgM antibodies with a Luminex–based quantification test that utilize beads coated with RBD, S1 domain of Spike and NP proteins respectively (Figure 2A).

**Figure 2.**
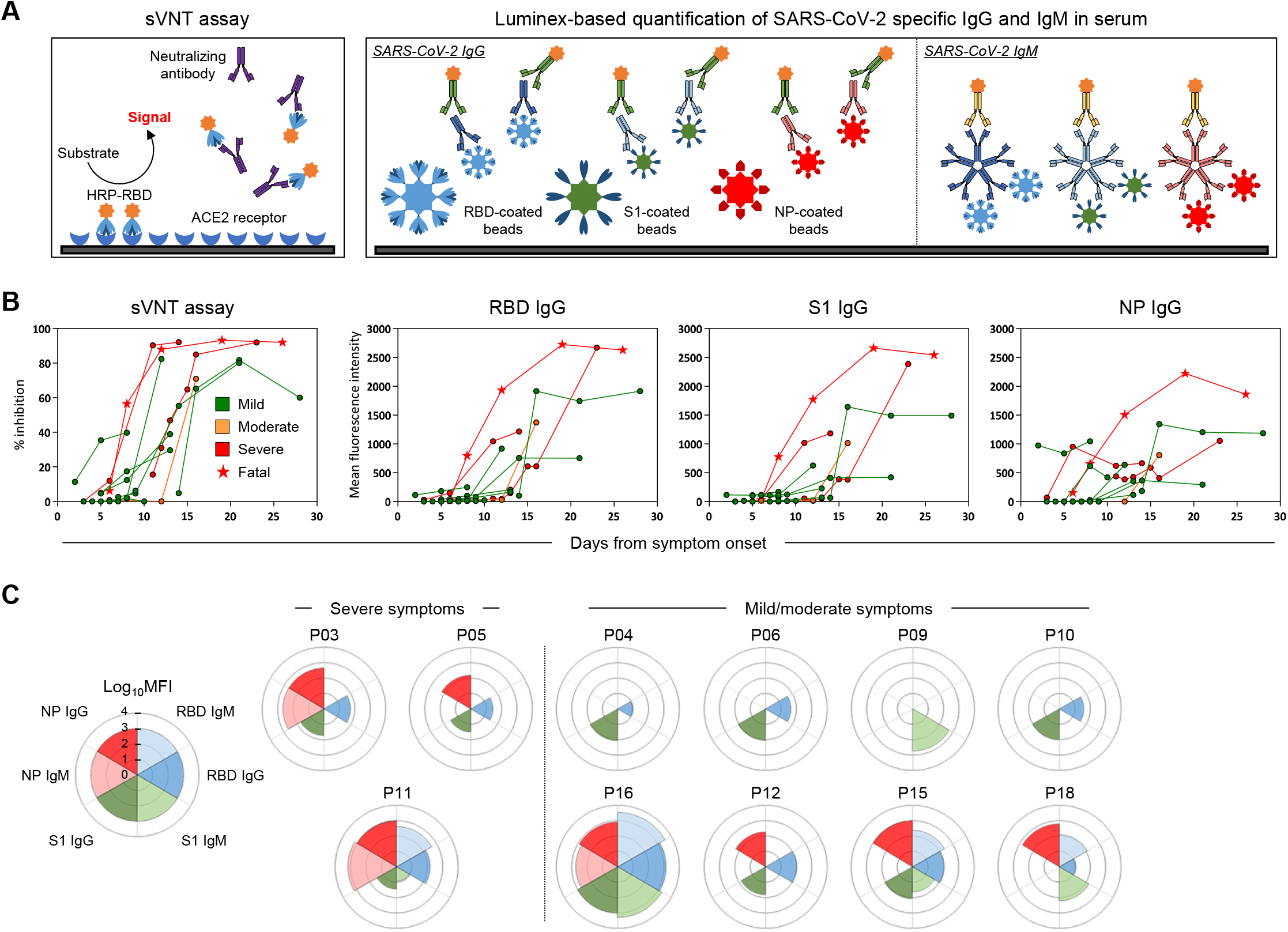
Longitudinal analysis of SARS-CoV-2 specific antibody-related responses in acute COVID-19 patients. A) Schematic representation of the surrogate virus neutralisation assay and the Luminex-based assay to quantify SARS-CoV-2 RBD-, S1- and NP-specific IgG and IgM antibodies. A cut-off to define significant virus neutralisation or antibody quantities was set at 20% inhibition for the sVNT assay (as defined in ref) and an MFI >100 for the Luminex based assay respectively. B) SARS-CoV-2 neutralisation and the relative quantities of specific IgG and IgM antibodies. C) Rose plots represent the quantity of RBD-, S1- and NP-specific IgG and IgM antibodies at the time of first detectable antibody response. Patient P02 has no detectable antibody response throughout and the corresponding rose plot is not shown.

Figure 2B shows that all COVID-19 patients developed neutralising antibodies with the exception of patient P02 who nonetheless cleared SARS-CoV-2 at day 13 post symptom onset (Figure 1). The peak neutralizing activity was achieved within 9-15 days post symptom onset. Of note, the two patients who first reached 90% virus neutralisation developed severe disease (P05-deceased and P03-severe). The kinetics of anti-RBD, anti-S1 and anti-NP IgG also peaked around the 10-20 days period, similar to previous observations (Isho et al., 2020; Ripperger et al., 2020)with higher peak levels of antibodies against NP and Spike detected in patient P05 who succumbed to the infection. Interestingly, when we analysed in parallel the kinetics of appearance of antibody responses against different proteins, we observed that patients with severe disease have an early NP-biased antibody response, while those with mild/moderate symptoms had either a spike-dominant or balanced response (Figure 2C). These kinetics of appearance is consistent with recent data (Atyeo et al., 2020; Sun et al., 2020) showing that an anti-nucleocapsid humoral response is preferentially induced over a Spike IgG response in severe COVID-19 patients.

### Dynamics of SARS-CoV-2 specific cellular responses

We next analysed the kinetics of SARS-CoV-2 specific T cell appearance during the RT-PCR positive phase of disease. Overlapping 15-mer peptide libraries covering the whole NP, Membrane (M), ORF7ab, ORF8, ORF3a, the NSP7 and NSP13 of ORF1ab and a pool of ~40 peptides containing all the confirmed T cell epitopes of Spike were used to stimulate PBMC in an IFN-γ ELISPOT assay (Figure 3A). Since samples of PCR+ patients could not be handled outside a BSL-3 laboratory, a more detailed immunological analysis performed by flow cytometry could only be performed after viral clearance (Supplementary Figure 1). During the initial phase of SARS-CoV-2 infection, the quantity of IFN-γ secreting cells after stimulation by the different peptides pools increased progressively with the peak of frequency detected within ~15 days after symptom onset in the majority of the tested patients (8 out of 12, Figure 3B and 3C) in line with a previous report (Weiskopf et al., 2020). In two patients, peak responses were detected beyond 20 days after symptom onset, while patient P05, who succumbed to disease before viral clearance, had no detectable IFN-γ secreting cells when stimulated with the different peptide pools until day 26 when stimulation with Spike peptides activated a weak response (Figure 3B). Importantly, large production of IFN-γ was detected at all time points in all patients after stimulation with PMA+Ionomycin, showing that global cellular functionality was not compromised (Figure 4A insert). The frequency of IFN-γ secreting cells reactive to all the different peptide pools was also quantified at least 1 month after resolution of infection. The magnitude of the IFN-γ response declined massively in 7 out of the 8 patients tested (Figure 3B), consistent with the waning of the cellular immune response that follows resolution of acute infection. The quantity and time of appearance of SARS-CoV-2 peptide-reactive cells was then analysed in relation to the virological and clinical parameters.

**Figure 3.**
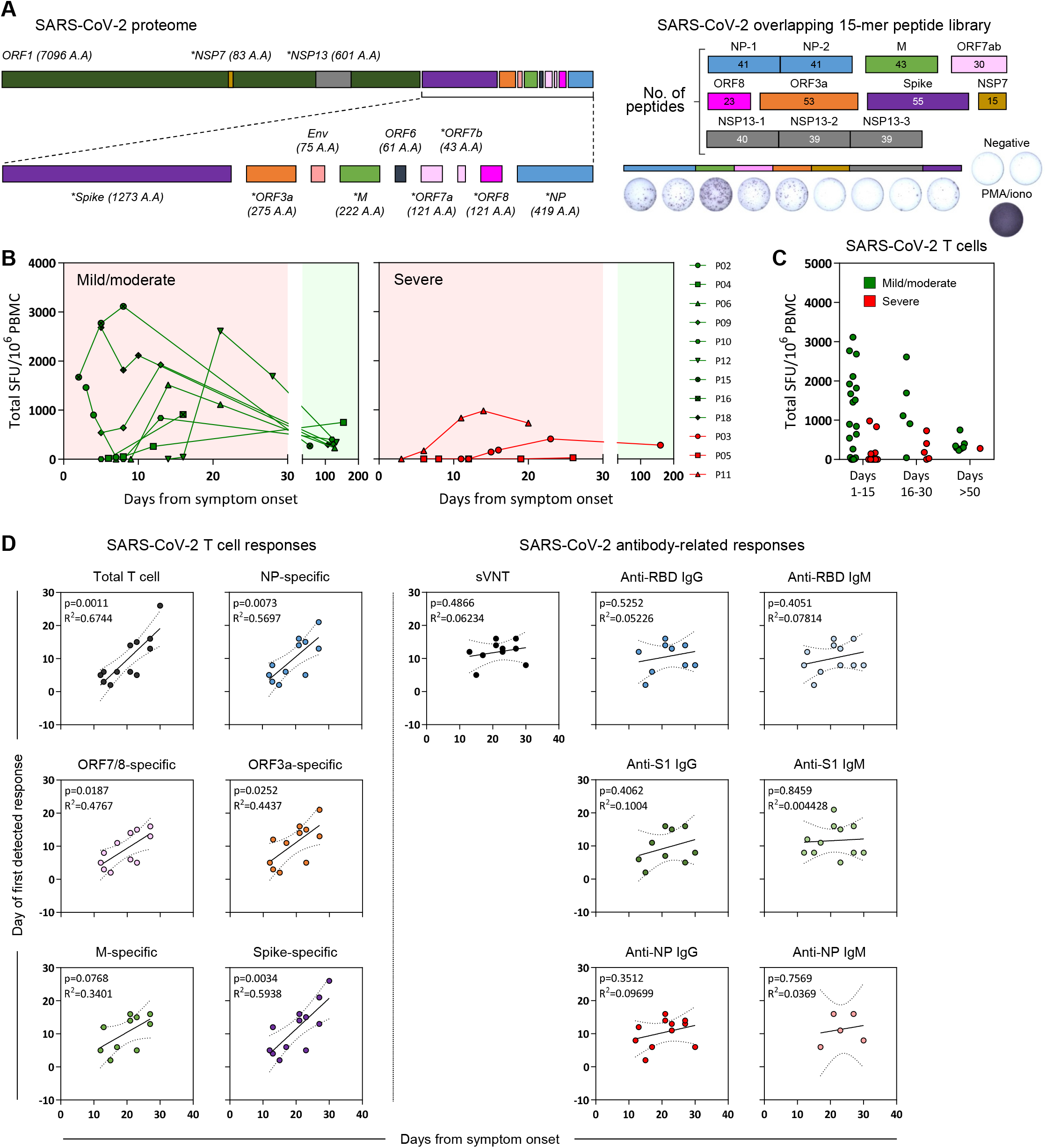
Longitudinal analysis of SARS-CoV-2 T responses in COVID-19 patients during acute infection and at convalescence. A) SARS-CoV-2 proteome organization; analysed proteins are marked by an asterisk. 15-mer peptides, which overlapped by 10 amino acids, comprising the nucleoprotein, membrane, ORF7ab, ORF8, ORF3a, NSP7 and NSP13 were grouped into 10 pools with the indicated number of peptides in each pool. 15-mer predicted peptides previously shown to activate Spike-specific CD8 and CD4 T cells were grouped into a single pool. PMA/ionomycin was used as a positive control for all samples analysed. B) Longitudinal analysis of the total SARS-CoV-2 T cell response in COVID-19 patients from onset of disease until convalescence. Individual lines represent single patients. C) Total SARS-CoV-2 T cell response detected in all COVID-19 patients (n=12) at days 1-15, 16-30 and >50 days after symptom onset. Patients with mild/moderate or severe symptoms are indicated. D) Correlation between the duration of infection and the number of days to the first detectable T cell response (Total, NP-specific, ORF7/8-specific, ORF3a-specific, M-specific or Spike-specific T cell response) or antibody-related response (sVNT, RBD-, S1- or NP-specific IgG and IgM) are shown in the respective dot plots. P-values and the corresponding r^2^ values are shown.

**Figure 4.**
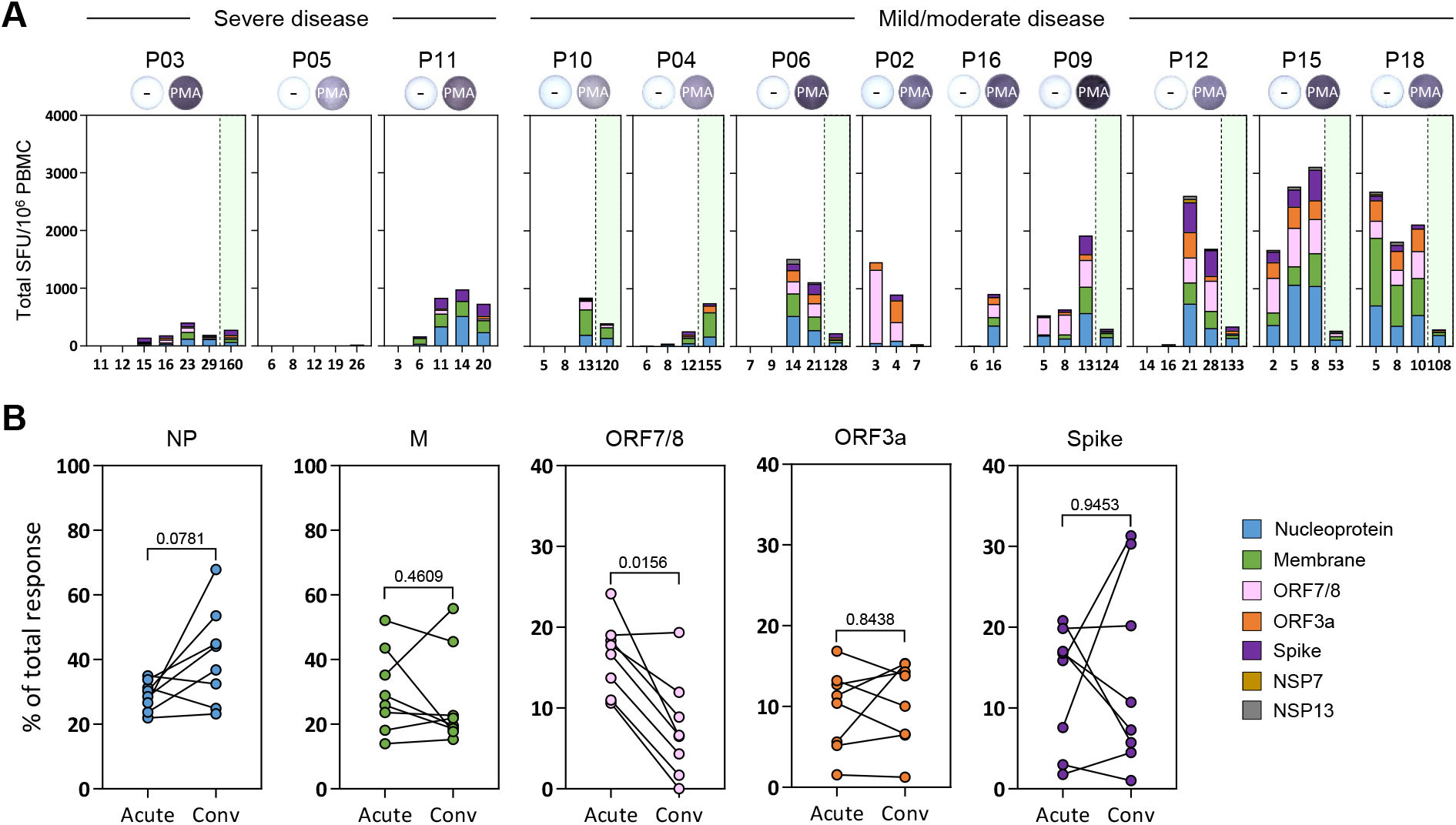
Hierarchy of cellular responses towards different SARS-CoV-2 proteins. A) Stacked bars denotes the frequency of peptide reactive cells in all COVID-19 patients against the indicated SARS-CoV-2 protein at all time points tested. Green shaded areas denote the convalescence phase of the disease. Positive controls are inserted for each patient. B) Plots show the proportion of peptide reactive cells attributed to the respective SARS-CoV-2 protein at the peak response during the acute phase and the convalescence phase of the disease (n=8). Wilcoxon matched-pairs test was used to evaluate the differences and the p-values are shown.

First, we observed that in contrast to the antibody quantity, the overall magnitude of SARS-CoV-2 peptide-reactive cells was not proportional to the severity of disease. Figure 2B shows that while higher quantities of IgG were observed in patients with severe compared to mild COVID-19, an opposite pattern was detected when the total IFN-γ response detected after stimulation by all the peptide pools was calculated. Higher frequencies of IFN-γ secreting cells both in the early (day 1-15) as well as late stages (day 15-30) were present in mild but not in the severe COVID-19 patients (Figure 3B and 3C). In addition, by analysing the association between the time of SARS-CoV-2 T cell appearance and the length of infection, we observed a statistically significant direct correlation between the early appearance of SARS-CoV-2 peptide-reactive cells (specific for NP, ORF7/8, ORF3a, M and Spike) and shorter duration of infection (Figure 3D). In contrast, no correlation was observed when we analysed the time of antibody and the length of infection. The temporal association of SARS-CoV-2 specific T cell appearance with reduced length of infection suggest that T cells play an essential role in the control of SARS-CoV-2 infection.

### Hierarchy of T cell immunogenicity towards different SARS-CoV-2 proteins

Finally, we performed a granular analysis of the ability of different SARS-CoV-2 proteins to stimulate IFN-y production in severe and mild cases of SARS-CoV-2 infection. Peptide pools covering all the different structural proteins (M, NP, ORF3a and Spike) and the ORF7/8 accessory proteins stimulate IFN-y production from the PBMC of all the acute patients with the exception of patient P05 (Figure 4A). In line with our previous report, NSP7 and NSP13 pools were rarely able to trigger IFN-y response in COVID-19 patients(Le Bert et al., 2020). In most of the acute patients, we observed the simultaneous presence of IFN-y producing cells specific for all the different peptide pools both during the acute phase of infection and at convalescence (Figure 4A). Of note, we observed that the ORF 7/8 peptide pool triggered a robust IFN-y response preferentially in the early phases of infection and only in patients with mild disease (Figure 3B and 4A). When we calculated the proportion of IFN-y producing cells triggered by different SARS-CoV-2 proteins at the time of first detection and at convalescence (>30 days after viral clearance), we observed that ORF7/8 responses waned almost completely at convalescence. In contrast, the proportion of cells stimulated by the peptide pools covering other SARS-CoV-2 proteins remained unchanged (M, ORF3a and Spike) or increased (NP) over time (Figure 4B). To demonstrate unequivocally that IFN-y secreting cells detected in our assays after peptide stimulation were indeed T cells, we performed a flow cytometry phenotypic analysis of IFN-y producing cells expanded after SARS-CoV-2 peptide stimulation of PBMCs. Unfortunately, such characterisations was performed only with PBMCs of SARS-CoV-2 patients at convalescence due to biosafety regulations that prevented us from analysing PBMCs collected during active infection (RT-PCR positive) outside a BSL-3 laboratory. As expected and already demonstrated, CD3+ T cells produced IFN-y after peptide pool stimulation (Supplementary Figure 1A). Most of the peptide responsive T cells were CD4 cells but CD8 T cells specific to NP peptide pools were also detected (Supplementary Figure 1B). We were also able to expand ORF7 or ORF8 specific T cells in 6 out of 7 patients in whom we detected, in the early phases of infection, a robust population of IFN-y producing cells after activation with the combined ORF7/8 peptide pools. Phenotypic analysis showed that these cells were all CD4 T cells. Thus even though we were unable to directly demonstrate that CD4 T cells were responsible for the IFN-y response triggered by ORF7/8 pools during the early phases of infection, such interpretation was strongly supported by the exclusive expansion of ORF7 and ORF8 specific CD4 T cells in the convalescents.

## Discussion

Despite the small number of patients analysed, our longitudinal analysis of the dynamics of virological and virus-specific immunological parameters during the acute phase of SARS-CoV-2 infection revealed a positive relation between induction of SARS-CoV-2 specific T cell responses and early control of infection. In addition, the quantity of virus-specific T cells present during the acute phase of infection was not proportional to the COVID-19 severity but was actually more robust in patients with mild disease. These data contrasted with the recent analysis of SARS-CoV-2 specific T cells in COVID-19 convalescent patients that reported a positive association of the frequency of SARS-CoV-2 specific T cells with disease severity(Peng et al., 2020). However in this work, the T cell response was measured in patients who were already in the convalescent phase. Since our longitudinal analysis showed that the frequency of SARS-CoV-2 specific T cells rapidly waned after SARS-CoV-2 clearance, the time of T cell analysis can significantly influence the magnitude of T cell response detected. The most robust T cell response was detected in patients with mild/moderate symptoms who cleared the virus early, while for example the patient with severe disease who succumbed to infection developed only a weak and monospecific T cell response detectable 26 days after symptom onset.

An opposite scenario was observed for antibodies. The two most severe COVID-19 patients studied here showed the most rapid and robust ability to achieve peak virus neutralisation and the overall quantities of SARS-CoV-2 specific antibodies were higher in severe than milder COVID-19 cases. These data confirm the numerous observations that have linked virus-specific antibody production (Hung et al., 2020; Long et al., 2020b) Ripperger:2020fq}(Ripperger et al., 2020) and B cell hyperactivation (Woodruff et al., 2020) with increased disease severity. Due to the significant link between early induction of T cells and shorter duration of the infection, the demonstration of the early induction of ORF7/8 specific cellular immunity can be of particular significance in viral control. For example, a 382 nucleotide deletion that truncates ORF7b and ORF8 leading to the elimination of ORF8 transcription had been reported at low frequencies in multiple countries (Su et al., 2020). While the mechanism leading to the acquisition of the genomic change is unresolved, the early induction of ORF7/8 specific T cells and a recent report of the robust early antibody response to ORF8 in SARS-CoV-2 infection would suggest that an immune-driven selection process could be involved (Hachim et al., 2020). Given that this viral variant was associated with a milder disease (Young et al., 2020), the early induction of ORF7/8 immunity is worthy of further investigation. It remains difficult to explain why ORF7/8 specific T cells were preferentially detected during the acute phase of infection. Recent findings show a corresponding increase in ORF8-specific antibodies during the early phases of SARS-CoV-2 infection (Hachim et al., 2020). However, there is no experimental evidences of a preferential early expression of ORF7/8 proteins in SARS-CoV-2 infected cells that might contribute to an increased immunogenicity of these accessory proteins in the early phases of SARS-CoV-2 infection. An alternative hypothesis is that pre-existing immunological memory to ORF7 or ORF8 might have caused a selective accelerated expansion of ORF7/8 T cells since ORF7/8 specific T cells can be occasionally detected in archived PBMC samples collected from healthy individuals before 2019 (Mateus et al., 2020). However, ORF7/8 is only expressed by SARS-CoV-1 and SARS-CoV-2 with little homology to other seasonal coronaviruses, hence the role of such peptide cross-reactive cells is puzzling and calls for a more detailed analysis of the effect of pre-existing immunity in the control or pathogenesis of SARS-CoV-2 infection and of the role of T cells specific for different antigens in SARS-CoV-2 protection.

## Acknowledgments

We want to express our gratitude to the study participants and personnel involved in ensuring the safety of the BSL3 laboratory operations.

## Author contributions

A.T.T., N.L.B. and A.B. designed the experiments. M.L., C.W.T, Y.Z. and G.J.S. performed the experiments in the BSL3 laboratory. W.N.C., C.W.T and L.W. performed the antibody analysis. K.K., C.T. and A.C. performed all other experiments. A.T.T., M.L., N.L.B. and A.B. analysed and interpreted the data. A.T.T. and A.B. prepared the figures and wrote the paper. B.Y., S.K., J.L. and D.L. recruited patients and provided all clinical data. A.B. designed, coordinated the study and provided funding.

## Methods

### Study design

Patients (n=12) were enrolled in this study after being admitted to the hospital and confirmed to be infected with SARS-CoV-2 based on a positive SARS-CoV-2 RT-PCR test as part of the PROTECT (National Healthcare Group Domain Specific Review Board reference number 2012/00917) and Novel Pathogens (CIRB ref. 2018/3045) studies. All participants provided written informed consent. Peripheral blood of acutely infected patients was collected in Mononuclear Cell Preparation tubes (CPT, BD Vacutainer) and transferred at 4°C to the biosafety level-3 (BSL3) facility for same-day processing. Blood from study participants at convalescent time points was obtained and processed in BSL2 laboratories. Six patients were male, six were female, their median age at time of admission was 52.5 years, ranging from 27 to 78 years. None of the patients received immunomodulatory treatments during the study period.

### Surrogate virus neutralization assay

sVNT assay to quantify the neutralising antibody response were performed as previously described (Tan et al., 2020). Briefly, sera from acutely infected patients were prepared in BSL3 containment and heat-inactivated prior to sVNT assay. HRP-conjugated RBD (Genscript) were pre-incubated with 1:20 diluted serum at 37°C for 1h, followed by addition to the Fc-chimeric human ACE2-coated MaxiSORP ELISA plate (Nunc) for an hour at room temperature. Colorimetric signal was developed using TMB substrate (KPL) after extensive PBST washes and the reaction was stopped with 1M HCl. Absorbance reading at 450 nm and 570 nm were obtained using Hidex Sense microplate reader (Hidex).

### Luminex analysis

SARS-CoV-2 RBD, S1 and N proteins (Genscript) were conjugated onto MagPlex microsphere (Luminex) using xMAP antibody coupling kit (Luminex). SARS-CoV-2 spike and N proteins specific antibodies were detected by pre-incubation of 100-fold diluted serum (in 1% BSA PBS) with conjugated microspheres (1250 beads/antigen) for 1h at room temperature, followed by 1:1000 diluted PE-conjugated anti-human IgG polyclonal antibody (eBioscience) or PE-conjugated anti-human IgM antibody for 1h at room temperature. The signal was detected using Luminex MAGPIX instrument.

### IFN-γ ELISPOT assay

15-mer peptides that spanned the entire ORF of genes eight SARS-CoV-2 proteins (NP, M, ORF7, ORF8, ORF3, S, NSP7, NSP13) and antibody responses to two SARS-CoV-2 proteins (NP, S) were synthesised with 10 amino acids overlap and were grouped into pools of approximately 40 peptides (Table S1). CPT tubes were centrifuged at 1500 rcf for 15 mins and approximately 2 ml of mononuclear cells located on top of the polyester gel were aliquoted and stored at −80°C. ELISPOT plates were prepared with IFN-γ coating antibody (MabTech) and peptides pools in 50 μl AIM-V medium (Gibco; Thermo Fisher Scientific) supplemented with 2% AB human serum (Gibco; Thermo Fisher Scientific) on the day of incubation with PBMCs. Vials containing PBMCs were thawed at RT, 2U Benzonase (SigmaAldrich) to remove remaining nucleic acids was added when fully thawed and incubated for 15 mins before centrifugation at 1000 rcf for 10 mins. The cell pellet was dissolved in 1 ml of AIM-V medium (Gibco; Thermo Fisher Scientific) supplemented with 2% AB human serum (Gibco; Thermo Fisher Scientific) before quantification at undiluted and 1:10 dilution using a Scepter 2.0 cell counter (Millipore). Approximately 2 x 10^5^ PBMCs in 100 μl per well were incubated in the presence of peptides overnight at 37°C and 5% CO2. After 24 hours, inoculum was removed and plates were washed six times with PBS. Biotinylated anti-IFN-γ antibody (MabTech) at a 1:2000 dilution in PBS/0.5% FCS was incubated at RT for two hours, followed by six wash steps with PBS and incubation of Streptavidin-AP (MabTech) at a 1:2000 dilution in PBS/0.5% FCS at RT for one hour. After another six PBS washes, 50μl of KLP BCIP/NBT phosphatase substrate (SeraCare) was added and incubated at RT in the dark for 5-15 mins. The reaction was stopped by washing the plate with water extensively when the chromogenic reaction produced clearly visible spots. Subsequently, the plates were allowed to air-dry and spot forming units (SFU) were analysed using an Immunospot reader and software (Cellular Technology).

### SARS-CoV-2 specific T cell lines

T cell lines were generated as follows: 20% of PBMC were pulsed with 10 μg/ml of the overlapping SARS-CoV-2 peptides (all pools combined) or single peptides for 1 hour at 37°C, subsequently washed, and co-cultured with the remaining cells in AIM-V medium (Gibco; Thermo Fisher Scientific) supplemented with 2% AB human serum (Gibco; Thermo Fisher Scientific). T cell lines were cultured for 10 days in the presence of 20 U/ml of recombinant IL-2 (R&D Systems).

### Flow cytometry analysis

Expanded T cell lines were stimulated for 5h at 37°C with or without SARS-CoV-2 peptide pools (2 μg/ml) in the presence of 10 μg/ml brefeldin A (Sigma-Aldrich). Cells were stained with the yellow LIVE/DEAD fixable dead cell stain kit (Invitrogen) and anti-CD3 (clone SK7; 3:50), anti-CD4 (clone SK3; 3:50), and anti-CD8 (clone SK1; 3:50) antibodies.. Cells were subsequently fixed and permeabilised using the Cytofix/Cytoperm kit (BD Biosciences-Pharmingen) and stained with anti-IFN-γ (clone 25723, R&D Systems; 1:25) and anti-TNF-α (clone MAb11; 1:25) antibodies and analysed on a BD-LSR II FACS Scan. Data were analysed by Kaluza (Beckman Coulter). Antibodies were purchased from BD Biosciences-Pharmingen unless otherwise stated.

### Statistical analysis

Regression analyses were performed using GraphPad Prism v7 software. Figures were prepared in Adobe Illustrator Creative Cloud.

## Supplementary Figure Legends

**Supplementary Figure 1.**
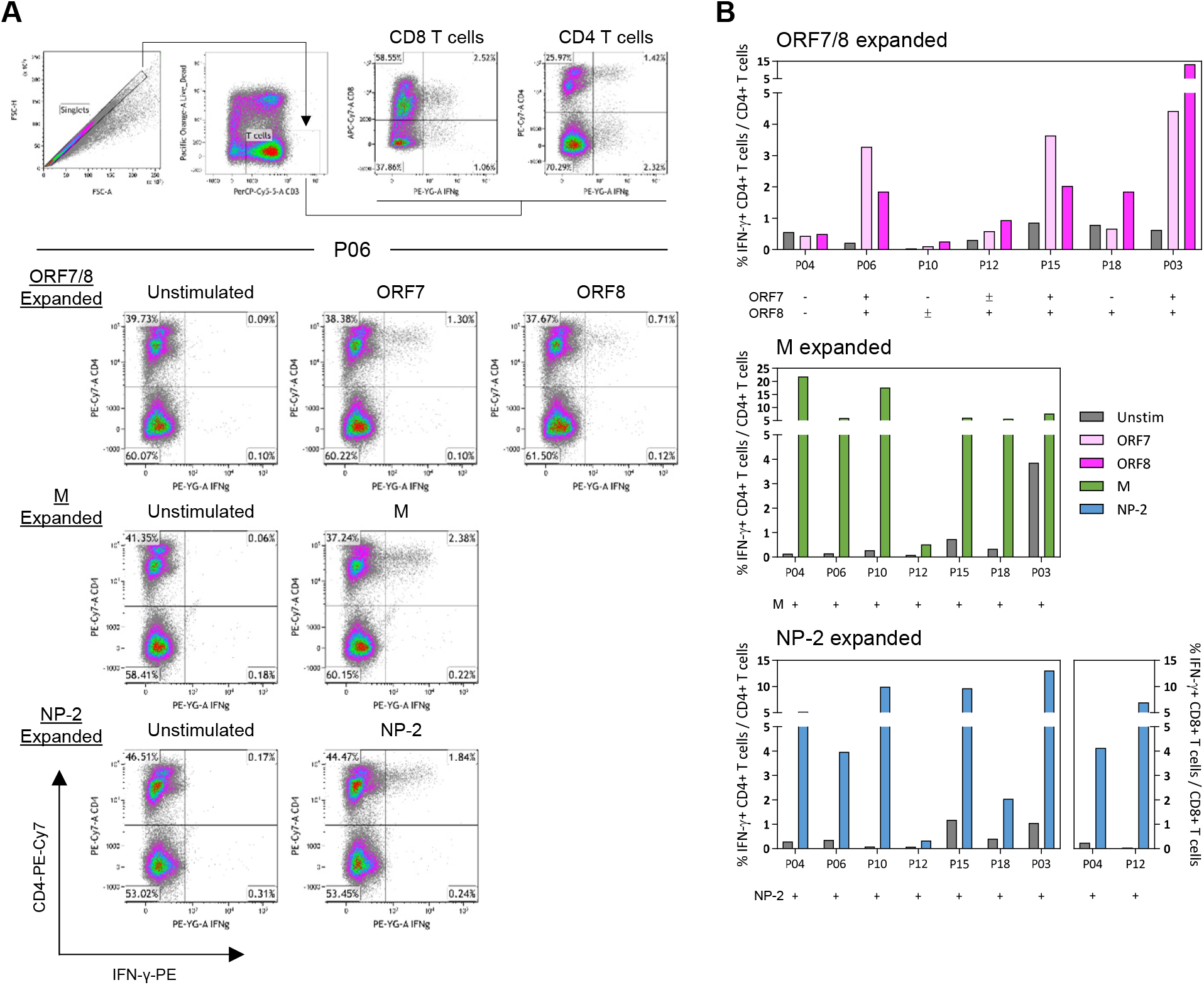
In vitro expanded SARS-CoV-2 specific T cell lines. A) Short-term T cell lines were generated using the respective SARS-CoV-2 peptide pools. Each line was then stimulated with the corresponding peptide pool used for expansion and the frequency of IFN-γ producing T cells was quantified. The flow cytometry gating strategy is shown above. Representative dot plots of ORF7/8, M and NP-2 specific T cell lines generated from patient P06 are displayed. C) Frequencies of IFN-γ producing CD4 or CD8 T cells of all in vitro expanded T cell lines generated from the convalescent PBMCs of respective COVID-19 patients (n=7).

## References

Atyeo, C., Fischinger, S., Zohar, T., Slein, M.D., Burke, J., Loos, C., McCulloch, D.J., Newman, K.L., Wolf, C., Yu, J., et al. (2020). Distinct Early Serological Signatures Track with SARS-CoV-2 Survival. Immunity 53, 524–532.e524.

Braun, J., Loyal, L., Frentsch, M., Wendisch, D., Georg, P., Kurth, F., Hippenstiel, S., Dingeldey, M., Kruse, B., Fauchere, F., et al. (2020). SARS-CoV-2-reactive T cells in healthy donors and patients with COVID-19. Nature 1–8.

Grifoni, A., Weiskopf, D., Ramirez, S.I., Mateus, J., Dan, J.M., Moderbacher, C.R., Rawlings, S.A., Sutherland, A., Premkumar, L., Jadi, R.S., et al. (2020). Targets of T Cell Responses to SARS-CoV-2 Coronavirus in Humans with COVID-19 Disease and Unexposed Individuals. Cell 181, 1489–1501.e15.

Gudbjartsson, D.F., Norddahl, G.L., Melsted, P., Gunnarsdottir, K., Holm, H., Eythorsson, E., Arnthorsson, A.O., Helgason, D., Bjarnadottir, K., Ingvarsson, R.F., et al. (2020). Humoral Immune Response to SARS-CoV-2 in Iceland. N Engl J Med NEJMoa 2026116–11.

Hachim, A., Kavian, N., Cohen, C.A., Chin, A.W.H., Chu, D.K.W., Mok, C.K.P., Tsang, O.T.Y., Yeung, Y.C., Perera, R.A.P.M., Poon, L.L.M., et al. (2020). ORF8 and ORF3b antibodies are accurate serological markers of early and late SARS-CoV-2 infection. Nat Immunol 20, 809–818.

Hung, I.F.-N., Cheng, V.C.-C., Li, X., Tam, A.R., Hung, D.L.-L., Chiu, K.H.-Y., Yip, C.C.-Y., Cai, J.-P., Ho, D.T.-Y., Wong, S.-C., et al. (2020). SARS-CoV-2 shedding and seroconversion among passengers quarantined after disembarking a cruise ship: a case series. The Lancet Infectious Diseases 20, 1051–1060.

Isho, B., Abe, K.T., Zuo, M., Jamal, A.J., Rathod, B., Wang, J.H., Li, Z., Chao, G., Rojas, O.L., Bang, Y.M., et al. (2020). Persistence of serum and saliva antibody responses to SARS-CoV-2 spike antigens in COVID-19 patients. Science Immunology 5.

Le Bert, N., Tan, A.T., Kunasegaran, K., Tham, C.Y.L., Hafezi, M., Chia, A., Chng, M.H.Y., Lin, M., Tan, N., Linster, M., et al. (2020). SARS-CoV-2-specific T cell immunity in cases of COVID-19 and SARS, and uninfected controls. Nature 584, 457–462.

Long, Q.-X., Liu, B.-Z., Deng, H.-J., Wu, G.-C., Deng, K., Chen, Y.-K., Liao, P., Qiu, J.-F., Lin, Y., Cai, X.-F., et al. (2020a). Antibody responses to SARS-CoV-2 in patients with COVID-19. Nat Med 26, 845–848.

Long, Q.-X., Tang, X.-J., Shi, Q.-L., Li, Q., Deng, H.-J., Yuan, J., Hu, J.-L., Xu, W., Zhang, Y., Lv, F.-J., et al. (2020b). Clinical and immunological assessment of asymptomatic SARS-CoV-2 infections. Nat Med 63, 706–712.

Mateus, J., Grifoni, A., Tarke, A., Sidney, J., Ramirez, S.I., Dan, J.M., Burger, Z.C., Rawlings, S.A., Smith, D.M., Phillips, E., et al. (2020). Selective and cross-reactive SARS-CoV-2 T cell epitopes in unexposed humans. Science eabd3871.

Peng, Y., Mentzer, A.J., Liu, G., Yao, X., Yin, Z., Dong, D., Dejnirattisai, W., Rostron, T., Supasa, P., Liu, C., et al. (2020). Broad and strong memory CD4+ and CD8+ T cells induced by SARS-CoV-2 in UK convalescent individuals following COVID-19. Nat. Immunol, 10.1038/s41590-020-0782-6.

Ripperger, T.J., Uhrlaub, J.L., Watanabe, M., Wong, R., Castaneda, Y., Pizzato, H.A., Thompson, M.R., Bradshaw, C., Weinkauf, C.C., Bime, C., et al. (2020). Orthogonal SARS-CoV-2 Serological Assays Enable Surveillance of Low Prevalence Communities and Reveal Durable Humoral Immunity. Immunity 1–49.

Rydyznski Moderbacher, C., Ramirez, S.I., Dan, J.M., Grifoni, A., Hastie, K.M., Weiskopf, D., Belanger, S., Abbott, R.K., Kim, C., Choi, J., et al. (2020). Antigen-Specific Adaptive Immunity to SARS-CoV-2 in Acute COVID-19 and Associations with Age and Disease Severity. Cell 10.1016/j.cell.2020.09.038.

Su, Y.C.F., Anderson, D.E., Young, B.E., Linster, M., Zhu, F., Jayakumar, J., Zhuang, Y., Kalimuddin, S., Low, J.G.H., Tan, C.W., et al. (2020). Discovery and Genomic Characterization of a 382-Nucleotide Deletion in ORF7b and ORF8 during the Early Evolution of SARS-CoV-2. mBio 11, 565.

Sun, B., Feng, Y., Mo, X., Zheng, P., Wang, Q., Li, P., Peng, P., Liu, X., Chen, Z., Huang, H., et al. (2020). Kinetics of SARS-CoV-2 specific IgM and IgG responses in COVID-19 patients. Emerg Microbes Infect 9, 940–948.

Takahashi, T., Ellingson, M.K., Wong, P., Israelow, B., Lucas, C., Klein, J., Silva, J., Mao, T., Oh, J.E., Tokuyama, M., et al. (2020). Sex differences in immune responses that underlie COVID-19 disease outcomes. Nature 1–9.

Tan, C.W., Chia, W.N., Qin, X., Liu, P., Chen, M.I.-C., Tiu, C., Hu, Z., Chen, V.C.-W., Young, B.E., Sia, W.R., et al. (2020). A SARS-CoV-2 surrogate virus neutralization test based on antibody-mediated blockage of ACE2-spike protein-protein interaction. Nat Biotechnol 395, 470–476.

Weiskopf, D., Schmitz, K.S., Raadsen, M.P., Grifoni, A., Okba, N.M.A., Endeman, H., van den Akker, J.P.C., Molenkamp, R., Koopmans, M.P.G., van Gorp, E.C.M., et al. (2020). Phenotype and kinetics of SARS-CoV-2-specific T cells in COVID-19 patients with acute respiratory distress syndrome. Science Immunology 5, eabd2071.

Woodruff, M.C., Ramonell, R.P., Nguyen, D.C., Cashman, K.S., Saini, A.S., Haddad, N.S., Ley, A.M., Kyu, S., Howell, J.C., Ozturk, T., et al. (2020). Extrafollicular B cell responses correlate with neutralizing antibodies and morbidity in COVID-19. Nat. Immunol. 11, 1708–1711.

Young, B.E., Fong, S.-W., Chan, Y.-H., Mak, T.M., Ang, L.W., Anderson, D.E., Lee, C.Y.-P., Amrun, S.N., Lee, B., Goh, Y.S., et al. (2020). Effects of a major deletion in the SARS-CoV-2 genome on the severity of infection and the inflammatory response: an observational cohort study. Lancet 396, 603–611.

Zhou, P., Yang, X.-L., Wang, X.-G., Hu, B., Zhang, L., Zhang, W., Si, H.-R., Zhu, Y., Li, B., Huang, C.-L., et al. (2020). A pneumonia outbreak associated with a new coronavirus of probable bat origin. Nature 579, 270–273.

